# Definition of a Small Core Transcriptional Circuit Regulated by AML1-ETO

**DOI:** 10.1101/2020.06.14.151159

**Authors:** Kristy R. Stengel, Jacob Ellis, Clare Spielman, Monica Bomber, Scott W. Hiebert

## Abstract

Transcription factors regulate gene networks controlling normal hematopoiesis and are frequently deregulated in acute myeloid leukemia (AML). Critical to our understanding of the mechanism of cellular transformation by oncogenic transcription factors is the ability to define their direct gene targets. While this seems to be a straight forward task, gene network cascades can change within minutes to hours, making it difficult to distinguish direct from secondary or compensatory transcriptional changes by traditional methodologies. We describe an approach utilizing CRISPR-based genome editing to insert a degron tag into the endogenous AML1-ETO locus of Kasumi-1 cells, as well as overexpression of a degradable AML1-ETO protein in CD34^+^ human cord blood cells, which is a an AML1-ETO-dependent pre-leukemia model. Upon addition of a small molecule proteolysis targeting chimera (PROTAC), the AML1-ETO protein was rapidly degraded in both systems. Furthermore, by combining rapid degradation with nascent transcript analysis (PRO-seq), RNA-seq and Cut&Run, we have defined the core AML1-ETO regulatory network, which consists of fewer than 100 direct gene targets. The ability of AML1-ETO to regulate this relatively small gene pool is critical for maintaining cells in a self-renewing state, and AML1-ETO degradation set off a cascade of transcriptional events resulting in myeloid differentiation.

## Introduction

Transcription factors are critical regulators of cell fate decisions and frequent targets of chromosomal translocations in a variety of human malignancies^1,2^. Given the importance of oncogenic transcription factors to the development and progression of human disease^3^, the ability to accurately define their direct gene targets is required to truly understand the underlying disease mechanisms, as well as uncover new nodes for therapeutic intervention. While efforts to define direct transcriptional networks have been underway for decades, numerous hurdles have presented themselves; however, the confluence of recent technological advances such as genome editing^4^, targeted protein degradation^5^, and nascent transcript analyses^6^ have left us poised to finally uncover the true functions of these critical oncogenes.

Traditionally, studies defining gene networks have relied on genetic inactivation of the transcription factor, either through gene deletion or RNAi approaches, which were then correlated with changes in steady-state mRNA levels. However, these approaches are limited in multiple ways^7^. First, the time required for genetic inactivation can take days, thus making it impossible to distinguish direct from indirect transcriptional events. Second, large differences in mRNA half-lives due to distinct post-transcriptional regulatory mechanisms can further cloud interpretations. These issues have been ameliorated to some degree by correlating the changes in gene expression with genome-localizing methods such as ChIP-seq or Cut&Run, yet DNA binding is not a good predictor of regulatory activity on nearby genes, and existing data sets often predict binding at many more loci than could possibly be regulated. However, by combining rapid and targeted transcription factor degradation with precision nuclear run-on sequencing (PRO-seq)^8^ to directly monitor gene transcription, we were able to identify high-confidence, direct transcriptional targets and eliminate the noise created by secondary transcriptional events. Here, we applied these methods to the analysis of the t(8;21) fusion protein, AML1-ETO (RUNX1-RUNX1T1; AE).

The t(8;21) chromosomal translocation is the single most frequent translocation in acute myeloid leukemia (AML)^9^. While this translocation is generally associated with a favorable prognosis following treatment with intense chemotherapy, approximately 40% of patients will eventually relapse^10^. Efforts to identify the direct transcriptional targets of AML1-ETO are critical to understanding the pathogenesis of this disease and ideally will lead to new therapeutic strategies. The AML1-ETO protein includes the DNA binding domain of the RUNX1 (AML1) transcription factor fused to almost the entirety of the ETO (RUNX1T1) transcriptional co-repressor^11^. Previous studies have sought to define the core AML1-ETO transcriptional program, which underlies its role in myeloid differentiation control^12,13^. However, these attempts have all assayed transcriptional changes days after knockdown of AML1-ETO expression, which led to the suggestion that AML1-ETO regulates hundreds to thousands of genes, yet many of these are likely secondary or compensatory changes in gene expression^14^. Furthermore, while many existing studies suggest that AML1-ETO primarily functions to silence the expression of critical RUNX1 target genes, numerous studies have also reported that AML1-ETO is required to maintain gene expression^15-20^. These seemingly conflicting studies suggest context-specific AE functions in transcriptional control, but also highlight the need for experimental approaches that are capable of unambiguously defining transcriptional targets and effects on RNA polymerase dynamics.

We utilized CRISPR-based genome editing to engineer the endogenous *AML1-ETO* locus to create a fusion with the FKBP12^F36V^ degron tag^21,22^. This allowed rapid degradation of the endogenous AML1-ETO fusion protein to define the molecular mechanism of transcriptional control genome-wide within the first 30 min to 2 hours of drug treatment. By coupling rapid protein degradation with PRO-seq to directly monitor changes in nascent transcription and Cut&Run to identify AML1-ETO–bound loci, we defined a surprisingly small core network of direct AML1-ETO-regulated genes. This core AML1-ETO transcriptional network is composed of roughly 60 genes, including critical mediators of myeloid differentiation and cell fate decisions such as the transcriptional regulators *CEBPA, PLZF* (*ZBTB16*), *NFE2* and *MTG16* (*ETO2* or *CBFA2T3*). Moreover, almost all AML1-ETO-controlled genes were up-regulated immediately following AML1-ETO degradation, suggesting that the fusion protein functions primarily as a transcriptional repressor.

Finally, we exogenously expressed AML1-ETO-FKBP12^F36V^ in CD34^+^ human cord blood stem and progenitor cells and found that in these primary cells, AML1-ETO again repressed a relatively small set of genes that were rapidly activated upon degradation of AML1-ETO-FKBP12^F36V^. Importantly, reactivation of these few gene targets was sufficient to induce differentiation in primary CD34^+^ cell cultures. Moreover, while the gene targets identified in Kasumi-1 cells and CD34^+^ HSPC cultures were largely overlapping, there were also differences. One notable difference was *GFI1b*, which was not up-regulated upon AE degradation in Kasumi-1 cells, yet was the second most up-regulated gene in CD34^+^ cell cultures. However, treatment of Kasumi-1 cells with an LSD1 inhibitor in combination with AE degradation cooperated to drive *GFI1b* re-expression as well as promote Kasumi-1 cell differentiation.

## Methods

### Cell Culture

Kasumi-1 cells were maintained in RPMI supplemented with 15% FBS, 50 U/ml penicillin, 50 μg/ml streptomycin, and 2mM L-glutamine. CD34^+^ human cord blood cells were obtained from StemCell Technologies, and long-term CD34^+^ HSPC cultures were established by overexpression of AML1-ETO or AML1-ETO-FKBP12^F36V^ as previously described^23^. Briefly, cells were thawed into pre-stimulation media (IMDM supplemented with 20% FBS, 55 μM BME, 50 U/ml penicillin, 50 μg/ml streptomycin, 2mM L-glutamine, and 100 ng/mL SCF, TPO, and Flt3-L). Following retroviral transduction, cultures were maintained in a myeloid culture media (IMDM supplemented with 20% FBS, 55 μM BME, 50 U/ml penicillin, 50 μg/ml streptomycin, 2mM L-glutamine, and 10 ng/mL SCF, Flt3-L, TPO, IL-3, and IL-6).

### dTAG-47

dTAG-47 was previously described^21^, and was synthesized by the Vanderbilt University School of Medicine Chemical Synthesis Core. dTAG-47 was reconstituted in DMSO and used throughout the study at a working concentration of 500 nM.

### CRISPR-Cas9

Kasumi-1 cells were engineered to incorporate a C-terminal FKBP12^F36V^-2xHA tag encoded in the endogenous AML1-ETO locus using CRISPR-Cas9-mediated gene editing. Briefly, crRNA targeting the 3’ end of *RUNX1T1 (ETO)-* TCTGAGTTCACGTCTAGCGA was annealed with tracrRNA (IDT). Annealed gRNAs were assembled into RNP complexes by incubation with Alt-R S.p. Cas9 Nuclease V3 (IDT) at room temperature. RNP complexes and equimolar HDR plasmid targeting the 3’ exon of *RUNX1T1* and containing *FKBP12*^*F36V*^*-2xHA-P2A-mCherry* were electroporated into Kasumi-1 cells. Successfully edited cells were sorted based on mCherry expression no sooner than 7 days following electroporation.

### Retrovirus production/transduction

Full length *AML1-ETO* was cloned into the *MSCV-IRES-GFP* retroviral expression vector. The indicated domain deletions and point mutations were generated by site-directed mutagenesis using the QuickChange Lightning Site-Directed Mutagenesis kit (Agilent). For CD34^+^ HSPC cultures, *AML1-ETO-FKBP12*^*F36V*^*-2xHA* or wild-type *AML1-ETO* were cloned into the *MSCV-IRES-mCherry* vector. For virus production, MSCV-based retroviral vectors + VSVG + *pMD-gag-pol* vectors were transfected into 293T cells with PEI. For infection of Kasumi-1 cells, 48 hours post-transfection, viral supernatant was filtered and spun onto cells in the presence of 8μg/mL polybrene. For infection of CD34^+^ cord blood cells, filtered viral supernatant was spun onto a retronectin-coated dish, cells were added to virus-coated plates in the presence of 8μg/mL polybrene and incubated in the presence of virus overnight at 37°C. Infected cells were washed the following day and placed in fresh media.

### Western Blotting

Cells were lysed in RIPA buffer supplemented with protease inhibitors and sonicated to shear DNA. Cleared lysates were resolved by SDS-PAGE, transferred to PVDF membrane and probed with specified antibodies. *Antibodies: α-*HA – Abcam ab18181, α-Lamin B-Santa Cruz sc-6217, α-IKAROS Santa Cruz sc-398265, α-GAPDH Santa Cruzsc-365062, α-Runx Homology Domain (RHD) made in house.

### Flow cytometry

Cells were washed and resuspended in at a concentration of 1 million cells per 100 μl in PBS + 0.5% BSA. Cells were stained with the indicated antibodies at the manufacturer’s suggested concentration for 20 minutes at 4°C, washed and acquired using a BD 4 Laser Fortessa flow cytometer.

### Antibodies

CD82 (BD, clone 423524), CD11b (BD, clone ICRF44), CD34 (BD, clone 581), CD38 (BD, clone HIT2).

### Q-RT-PCR

Kasumi-1 AE-FKBP cells were treated for 3 days with 500 nM dTAG-47 alone, 100 nM LSD1i (GSK2879552, Selleckchem, S7796) alone, dTAG and LSD1i, or dTAG, LSD1i and 1 μM ATRA (Sigma-Aldrich, R2625). Total RNA was isolated by Trizol extraction according to the manufacturer’s protocol. 2 μg of RNA was used for reverse transcription (High Capacity cDNA Reverse Transcription Kit, ThermoFisher). Quantitative PCR detecting *Gfi1b* and *Actb* was carried out using SYBR green master mix (Bio-Rad). PCR primers: *Gfi1b* F-AGAAGGCTCACACCTACCAC, *Gfi1b* R-GCTAGGCTTGTAGAATGGGGG *Actb* F-ACCTTCTACAATGAGCTGCG, *Actb* R-CCTGGATAGCAACGTACATGG.

### Methylcellulose assays

Kasumi-1-AE-wild-type or Kasumi-1-AE-FKBP cells were treated for 24 hours with 500 nM dTAG-47 prior to plating in methylcellulose (MethoCult H4434, StemCell Technologies). 500 cells were plated in 1.1 mL Methocult supplemented with 500 nM dTAG-47. Colonies were counted 12 days after plating, cells harvested and re-plated in Methocult with fresh dTAG molecule.

### RNA-seq

All RNA-seq experiments were performed in biological replicates. Total cellular RNA was isolated using Trizol Reagent (Invitrogen) according to the manufacturer’s protocol. 5 μg of RNA was subject to DNase digestion, phenol-choloroform extracted and ethanol precipitated. RNA was submitted to Vanderbilt University Medical Center VANTAGE Core for poly-A enrichment-based library preparation and sequencing on the Illumina NovaSeq (PE-100). *Data Analysis.* Pre-processed reads were aligned to the human genome (hg19, downloaded from UCSC) using TopHat (v2.0.11) and differential gene expression was determined using Cuffdiff (v.2.1.1) as previously described^24^.

### PRO-seq

All PRO-seq experiments were performed in biological replicates. Nuclear run-on and PRO-Seq library construction were performed as previously described^25^, with *in vitro* transcribed and biotinylated *Luciferase, GFP*, and *NeoR* transcripts included as spike-in controls. Briefly, cells were treated with 500 nM dTAG-47 for 30 min, 1 hour, 2 hour, 6 hour or treated with 500 nM dTAG-47 alone, 100 nM GSK2879552 alone or dTAG + GSK2879552 for 24 hours prior to nuclei isolation. 30×10^6^ nuclei were used per run-on. Nuclear run-ons were carried out in the presence of 375 μM of biotin-11-CTP, ATP, UTP and GTP and 0.5% sarkosyl for 3 minutes at 30°C. Total RNA was hydrolyzed with 0.2N NaOH and nascent RNAs that had incorporated biotin-11-CTP were isolated by streptavidin bead binding. Prepared libraries were PAGE purified and provided to Vanderbilt University Medical Center VANTAGE Core for sequencing on an Illumina Nextseq 500 (SR-75). *Data Analysis.* Adaptors were trimmed and reads shorter than 15 bp were removed using Trimmomatic-0.32^26^. Reverse complement sequences were obtained using FASTX toolkit (v 0.0.13) prior to aligning to the human genome (hg19) using Bowtie2 (v2.2.2)^27^. Analysis of PRO-seq data was carried out using the Nascent RNA Sequencing Analysis (NRSA) pipeline^28^.

### Cut&Run

Cut&Run was performed with anti-HA (Cell Signaling Technology, C29F4) and anti-Rabbit secondary antibody (Invitrogen, 31238). Briefly, 250,000 cells were washed in buffer containing digitonin (0.01% digitonin for Kasumi-1 cells, 0.04% digitonin for CD34^+^ HSPCs) and bound to 10 μl of activated Concanavalin A beads (Bangs Laboratories Inc., BP531). Bead-bound cells were incubated with anti-HA (1:800) overnight, washed, and incubated with anti-rabbit secondary (1:100) for 1 hour. After washing, cells were incubated with CUTANA pA/G-MNase (Epicypher) and targeted chromatin digestion initiated by the addition of 100 mM CaCl_2_ and allowed to proceed for 2 hours at 4°C, at which time stop buffer containing *S.cerevisiae* spike-in DNA was added. Released chromatin fragments were purified by phenol-chloroform extraction followed by ethanol precipitation. Libraries were generated using the NEBNext Ultra II DNA Library Prep Kit and sequenced on the Illumina NovaSeq (PE-100) at Vanderbilt University Medical Center VANTAGE core. *Data Analysis.* Adaptors were trimmed with Trimmomatic-0.32 prior to aligning to the hg19 genome with Bowtie2 (v 2.2.2). Spike-in reads were removed and quantified. Peaks were called using MACS2 peak caller (narrowPeak; q-0.001; v 2.0.10.20131216)^29^. Peaks with significantly lower signal following dTAG treatment were identified with DiffBind^30^ and subsequent data analysis performed using Homer^31^ and deepTools (v 3.4.3)^32^.

### ChIP-seq

ChIP-seq was performed using anti-H3K27ac (abcam ab4729) on 10 million Kasumi-1 AE-FKBP cells at 0- and 24-hours post-treatment with dTAG-47 and with *Drosophila* S2 cell spike-in. Cells were cross-linked with 1% formaldehyde for 10 minutes and quenched with 125 mM Glycine. Following cell lysis, chromatin was sonicated with a Biorupter (Diagenode) to generate 300-600 bp chromatin fragments and immunoprecipitated with antibody plus Protein A:G beads. Library construction was carried out using the NEBNext Ultra II DNA Library Prep Kit and sequenced on the Illumina NovaSeq (PE-100) at Vanderbilt University Medical Center VANTAGE core. *Data Analysis:* Adaptors were trimmed with Trimmomatic-0.32 prior to aligning to the hg19 genome with Bowtie2 (v 2.2.2). Spike-in reads were removed and quantified. Bigwig files were generated using deepTools and normalized based on spike-in reads.

## Results

### CRISPR-mediated “dTAGging” of endogenous AML1-ETO

Degron tags provide the opportunity to rapidly remove *endogenous* proteins and assess the direct changes in gene expression to define the molecular mechanism of transcription factor action. Most importantly, this system avoids compensation or secondary effects in transcription networks that can occur in just a few hours. We used CRISPR/Cas9 to insert the FK506 binding protein, *FKBP12*^*F36V*^ just prior to the stop codon in the endogenous allele of *AML1-ETO* in Kasumi-1 (Fig. 1A). This created a fusion protein containing all of AML1-ETO fused to FKBP12^F36V^ along with a double HA epitope tag. Successfully edited Kasumi-1 cells were isolated by fluorescent activated cell sorting (FACS) using co-expressed mCherry^21^. Western blot analysis detected a mobility shift consistent with the addition of FKPB12 and this slow migrating AML1-ETO was also detected by anti-HA (Fig. 1B). The FKBP12^F36V^ tag rendered AML1-ETO sensitive to the heterobifunctional molecule, dTAG-47, which joins the synthetic FKBP12^F36V^ ligand to a lenalidomide-derivative that recruits the Cereblon E3 ubiquitin ligase (CRBN)^22^. Thus, treatment of Kasumi-1 *AML1-ETO-FKBP* cells with 500 nM dTAG-47 caused rapid degradation, with the majority of the protein lost within the first 1-2 hr (Fig. 1C). Moreover, this affect appeared specific to AML1-ETO, as protein levels of the known lenalidomide/CRBN target, IKAROS^33,34^, were unaffected (Fig. 1C).

**Figure 1.**
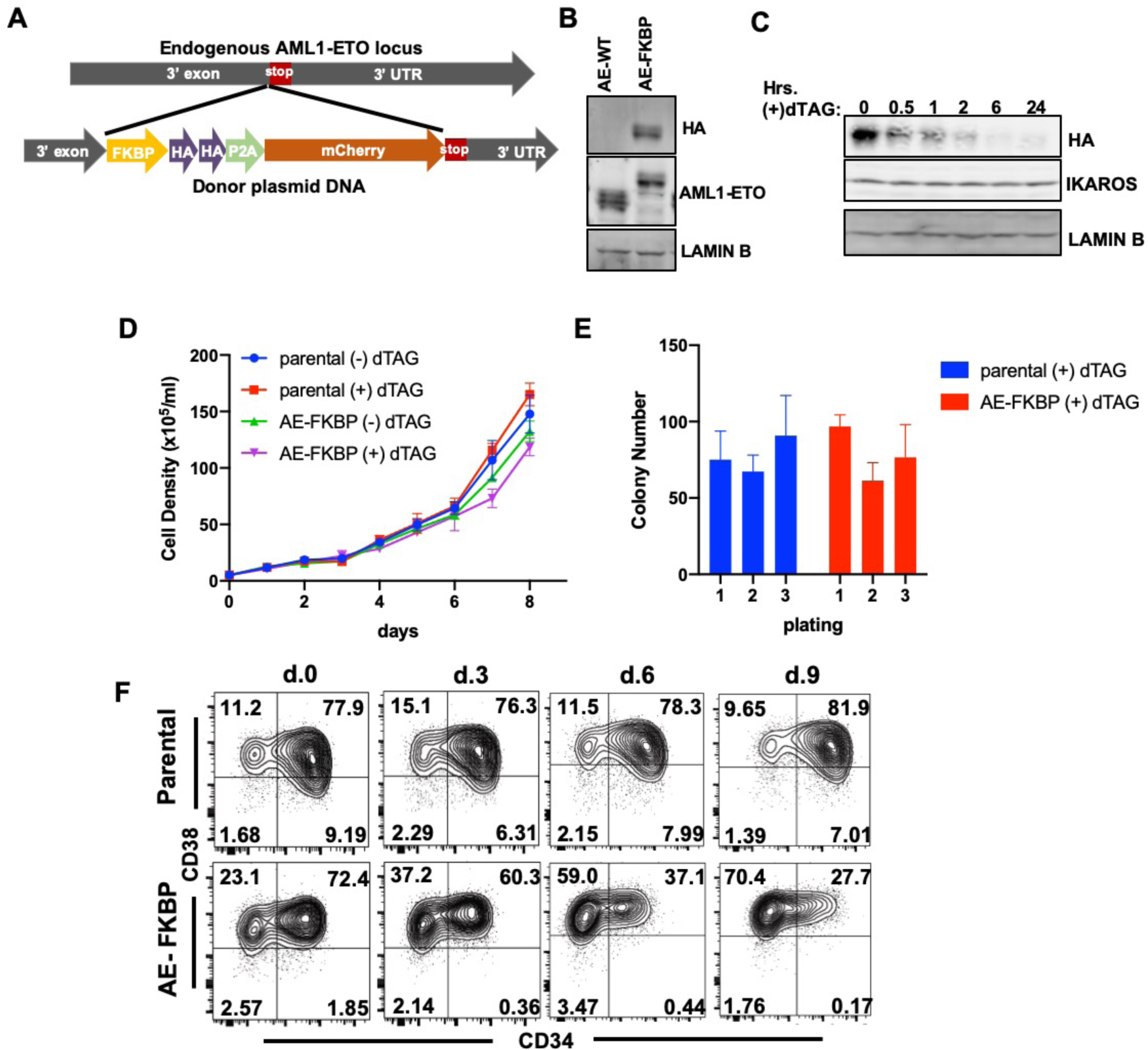
Targeting of an FKBP12-F36V degron tag to the endogenous AML1-ETO locus allows protein degradation upon dTAG-47 treatment. (A) Schematic depicting CRISPR-HDR based targeting approach to insert an FKBP12^F36V^ degron tag and 2xHA tag into the 3’ end of the endogenous AML1-ETO (AE) locus of the Kasumi-1 t(8;21) cell line. (B) Western blot from lysates of parental Kasumi-1 cells or AML1-ETO-FKBP12^F36V^-2xHA (AE-FKBP) Kasumi-1 cells. Anti-HA identifies the AE-FKBP protein, while an antibody recognizes the Runx homology domain identifies both wild-type AE and the larger AE-FKBP protein. (C) Kasumi-1-AE-FKBP cells were treated with 500 nM dTAG-47 to induce AE-FKBP degradation for the indicated times. AE-FKBP protein levels were monitored by blotting with anti-HA. IKAROS, a known CRBN E3 ligase target, was unaffected by dTAG-47 treatment. (D) Parental Kasumi-1 cells or Kasumi-1-AE-FKBP^F36V^ cells were treated with 500 nM dTAG-47 for 8 days and growth monitored by cell counts. (E) Parental Kasumi-1 cells or Kasumi-1-AE-FKBP^F36V^ cells were cultured in the presence of 500 nM dTAG-47 for 24 hours and then plated in methylcellulose in the presence of 500 nM dTAG-47. Colonies were counted and replated every 12 days. Parental Kasumi-1 cells or AE-FKBP^F36V^-expressing cells were treated with 500 nM dTAG-47. Cells were analyzed for surface CD34 and CD38 expression by flow cytometry every three days for 9 days. At each timepoint, cells were washed and reseeded in fresh media containing 500 nM dTAG-47.

AML1-ETO expression caused leukemia in mouse models, indicating that it is not only an oncogene^35-37^, but that it should be a good therapeutic target. In addition, shRNAs to ETO inhibited the growth of Kasumi-1 and SKNO-1 cells, but the effects were modest or transient in most instances^16,38,39^. We found that specific removal of the fusion protein from Kasumi-1 cells had only modest effects with no statistical differences on cell growth (Fig. 1D) or colony formation/replating assays in methylcellulose (Fig. 1E). However, degradation of AML1-ETO caused loss of the CD34^+^ cell population (Fig. 1F), which suggested that the fusion protein was regulating differentiation rather than directly regulating cell growth and/or viability. Importantly, while degradation of AE eliminated the CD34^+^ population thought to contain the leukemia stem cell, degradation of AE alone was not sufficient to fully trigger terminal myeloid differentiation.

### Defining the core AML1-ETO transcription signature

Precision nuclear run-on sequencing (PRO-seq) has been effectively employed at short intervals after small molecule treatment (e.g. BET inhibitors^25^, CDK9 inhibitors^40^, and CDK7 inhibitors^41^) to discern the mechanism of action of key transcriptional regulators. Therefore, we performed RNA-seq and PRO-seq at short intervals after dTAG-47 treatment to identify the early, likely direct, effects of AML1-ETO loss on gene expression. RNA-seq analysis showed a wave of changes in the steady state mRNA pools beginning at 2hr after addition of the dTAG and by 4hr distinct changes were apparent (Fig. 2A). By 8hr after addition of dTAG-47, some of the genes activated at 2hr and 4hr were already beginning to wane, and new sets of genes were activated. A similar trend was observed at 24hr with the genes most intensely activated at this late time point having shown minimal activation at 2-4 hr. It is also important to note that even though AML1-ETO is commonly regarded as a transcriptional repressor, it has been reported to activate many genes^15-20^, and RNA-seq identified roughly as many mRNAs that decreased as increased upon AML1-ETO degradation (Fig. 2B).

**Figure 2.**
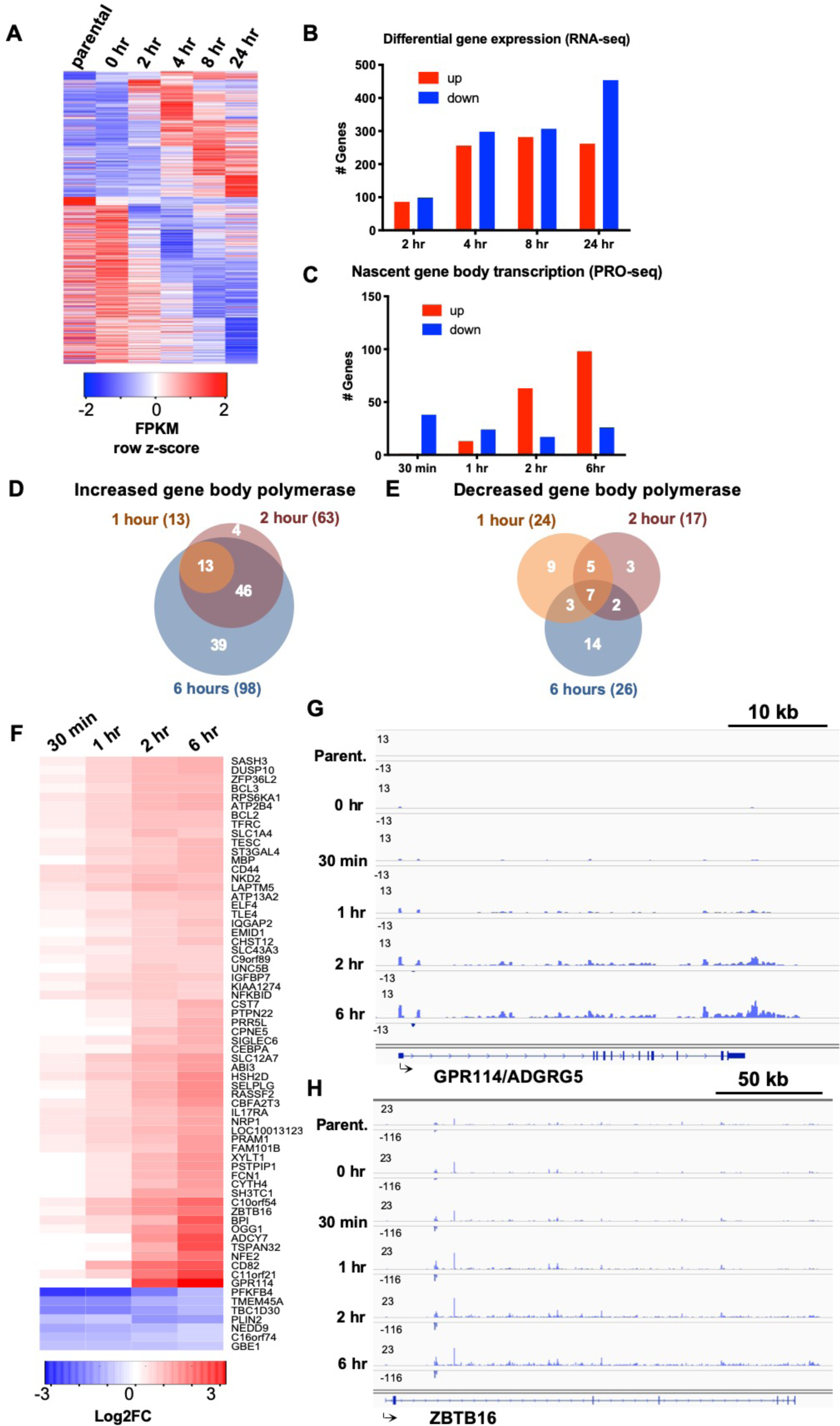
AML1-ETO represses the expression of fewer than one-hundred genes. (A) RNA-seq was performed on parental Kasumi-1 cells or Kasumi-1-AE-FKBP^F36V^ cells at the indicated timepoints following dTAG treatment (B) Quantitation of the number of genes identified as significantly up-regulated (up > 1.5-fold; q<0.05) or down-regulated (down > 1.5-fold; q<0.05) by RNA-seq at the indicated timepoints. (C) Quantitation of the number of genes with increases (up > 1.5-fold; q<0.05) or decreases (down > 1.5-fold; q<0.05) in gene transcription as identified by PRO-seq at the indicated timepoints. (D) Venn diagram depicts the overlap of genes exhibiting increased transcription at 1, 2, and 6 hours following AE degradation. (E) Venn diagram depicts the overlap of genes exhibiting decreased transcription at 1, 2, and 6 hours following AE degradation. (F) Heatmap showing the change in gene transcription of 59 up-regulated genes and 7 down-regulated genes identified by PRO-seq over time compared with 0 hr controls. Screenshots illustrate changes in PRO-seq signal at the *GPR114/ADGRG5* (G) and *ZBTB16* (H) loci over time.

A huge advantage of PRO-seq is that by examining nascent transcripts, changes in polymerase dynamics were detected within 30 min of addition of many compounds that target transcription^25,41^. Remarkably, examining nascent transcription detected the loss of a small cohort of genes within the first hour of dTAG-47 addition to the cultures (Fig. 2C). However, it is notable that the genes whose transcription decreased were not sustained across the time course, such that only 7 genes were consistently down-regulated at 1, 2, and 6hr (Fig. 2E). That is, genes that might have been “activated” by AML1-ETO were not consistent between time points (e.g. Fig. 2A, right panel). By contrast, the 13 genes that were induced 1.5-fold (using a 50 bp sliding window across the gene body^28,42^) at 1hr became more intense at 2hr, and 59 genes induced at 2hr were still induced at 6hr with a total of 98 genes induced in the first 6hr (Fig. 2D, and 2F). Thus, the cohort of genes that were activated upon AML1-ETO degradation were consistent and grew in intensity over time (Fig. 2C, 2D and 2F). This 59 gene signature included the canonical AML1-ETO-regulated genes *CEBPA, CD82, ETO2* and *ZBTB16* (Fig. 2F) that were induced at least 1.5-fold within 2hr (FDR-adjusted p value: q < 0.05). *PU.1* also showed a trend toward activation, but because its levels are only reduced by roughly 50% by AML1-ETO^43^, the induction did not meet our strict statistical cutoffs. Gene tracks showing transcriptionally engaged polymerase across the *ZBTB16* (*PLZF*) and *GPR114* show an observable increase in transcription at these loci by 1 hour, with an even more dramatic increase observed by 6 hours following AML1-ETO degradation (Fig. 2G, 2H).

Heatmaps constructed around the transcritional start sites of genes significantly activated or repressed by the degradation of AML1-ETO at all timepoints showed no effects on RNA polymerase pausing or elongation (Supplemental Fig. 1A). Furthermore, histograms depicting the PRO-seq signal around the TSS of genes activated upon AML1-ETO degradation revealed an increase in promoter proximal polymerase that slightly preceded the observed increase in gene body polymerase, suggesting an increase in transcription initiation upon AML1-ETO degradation led to increased gene body transcription (Supplemental Fig. 1B). We also intersected the PRO-seq and RNA-seq data, which showed that by 4-8hr the RNA-seq analysis had captured the same gene signature as the PRO-seq analysis, but also identified a number of changes in steady state transcripts that lacked a clear transcriptional basis (Supplemental figure 1C and 1D).

### Functional annotation of AML1-ETO binding sites

Given that the targeted degradation of AML1-ETO identified a much smaller core transcriptional signature as compared to genetic deletion or RNAi approaches, we performed CUT&RUN analysis to determine if the FKBP-2xHA tag had altered the AML1-ETO target selection (Fig. 1A). Because CUT&RUN does not use formaldehyde crosslinking, this also provides an analysis of native DNA binding by the endogenous fusion protein. This approach identified over 30,000 peaks that were significantly decreased upon degradation of the fusion protein (Supplemental Fig. 2A and 2B). The vast majority of these peaks overlapped with previous ChIP-seq data^44^ as well as Cut&Run performed on parental Kasumi-1 cells using anti-ETO (Supplemental Fig. 2C). Indeed, when the published ChIP-seq data and our Cut&Run were plotted around all the center of the peaks, the signals were virtually indistinguishable (Supplemental Fig. 2C). Thus, both wild-type AML1-ETO (anti-ETO) and AML1-ETO-FKBP12^F36V^-2xHA (anti-HA) genome binding was highly consistent with previously published AML1-ETO ChIP-seq data sets.

By ranking the AML1-ETO peaks based on the intensity of the Cut&Run signal (Fig. 3A), the peaks were placed into one of three clusters with the most intense peaks representing only the top 1989 peaks (Fig. 3A). We used HOMER to annotate peaks to the nearest gene, and found that AML1-ETO peaks within clusters 1 and 2 showed an even split between promoter and enhancer (intronic and intergenic) binding, while cluster 3 showed predominantly intronic and intergenic peaks. To functionally annotate peaks, we performed ChIP-seq for H3K27ac, a mark of active enhancers and promoters, before and after degradation of AML1-ETO with dTAG-47. By plotting this signal around all peaks, even when separated into clusters 1-3, we found that there was no increase in H3K27ac following AE degradation. In fact, if anything the H3K27ac signal was slightly decreased following dTAG treatment (Fig. 3C, upper). However, based on nearest gene annotation, the AML1-ETO peaks that were associated with the 59 gene repression signature identified by PRO-seq showed a robust increase in H3K27 acetylation upon AML1-ETO-degradation (Fig. Fig. 3C, lower). Overall, these peaks represented roughly 1% of the total HA-AML1-ETO peaks detected (Supplemental Fig. 3A), and were more highly enriched in clusters 1 & 2 than cluster 3 (Fig. 3C, lower). Thus, the major changes in acetylation occured in clusters 1 and 2, and not cluster 3, which were weaker binding sites.

**Figure 3.**
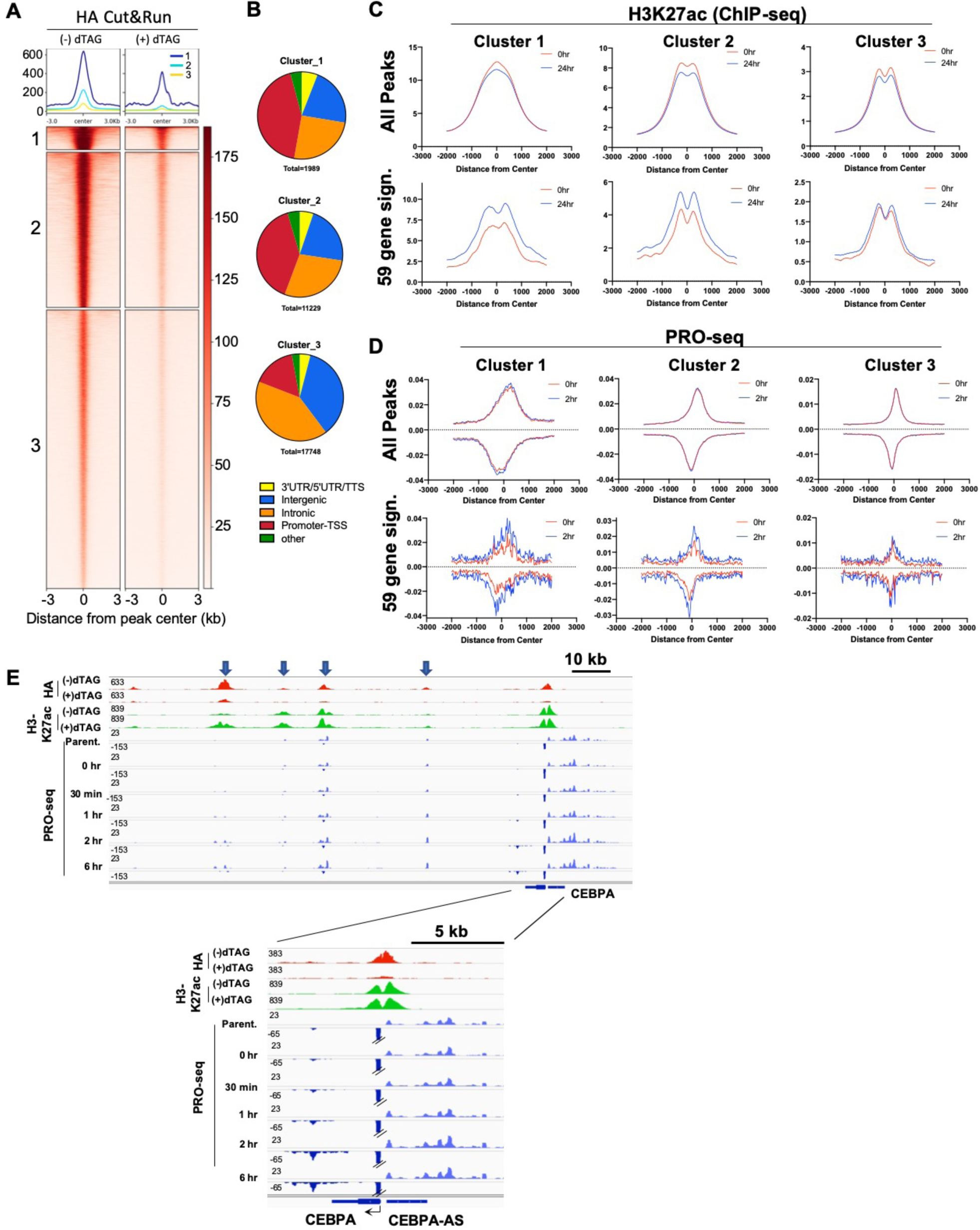
Intersection of AE binding with PRO-seq and H3K27ac ChIP-seq defines high-confidence AML1-ETO targets. (A) Cut&Run analysis was performed using anti-HA to detect AE-FKBP^F36V^-2xHA binding before and after 18-hour treatment with 500 nM dTAG-47. Peaks identified as significantly changed after dTAG-treatment by DiffBind are depicted in the heatmap. K-means clustering was used to segment peaks based on signal intensity (Clusters 1-3). (B) Peaks were annotated using Homer’s annotatePeak.pl to identify the nearest neighboring gene and the relative location of binding sites for each cluster identified. (C) Histograms depict the change in H3K27ac signal around HA-peaks following a 24-hour dTAG treatment. The H3K27ac ChIP-seq signal was plotted for +/- 2kb around all HA-peak centers for each cluster (upper) or plotted only for peaks associated with the 59 gene AE repression signature identified in Figure 2 (lower). (D) PRO-seq signal at 0 hours and 2 hours following dTAG treatment is plotted around all HA-peak centers in each cluster (upper) or around only those peaks associated with the 59 gene AE repression signature. (E) Screenshots depicting HA (AML1-ETO) Cut&Run, H3K27ac ChIP-seq and enhancer and gene body transcription (PRO-seq) at the *CEBPA* locus. H3K27ac was detected by ChIP-seq at 24 hours following AE-degradation by dTAG. PRO-seq was carried out at the indicated timepoints. Arrows indicate intergenic enhancers

Like gene promoters, active enhancers are characterized by bi-directional transcription inititaion. Therefore, we also plotted the PRO-seq signal +/-2,000 bp around identified AML1-ETO-binding sites. Much like H3K27ac, there was no change in active polymerase around AML1-ETO binding sites when all peaks are included in the analysis (Figure 3D, upper). However, when we focused on peaks associated with the 59 gene repression signature, we observed an increase in PRO-seq signal 2 hours following AE degradation at genes associated with clusters 1 & 2 (Fig. 3D, lower). Thus, it appears that functional AML1-ETO peaks are enriched in clusters 1 and 2 and characterized by both an increase in H3K27ac and active polymerase following AE degradation. For instance, the *CEPBA* locus is characterized by at least four downstream enhancers. The most distal enhancer had the most prominent AML1-ETO (HA) peak that was lost following dTAG treatment, and correlates with both an increase in H3K27ac and an increase in active polymerase within 6 hours (Figure 4E, upper). Moreover, all of these changes were associated with an observeable increase in *CEBPA* gene transcription by 1 hour that becomes even more significant by 6 hours (Figure 4E, lower).

**Figure 4.**
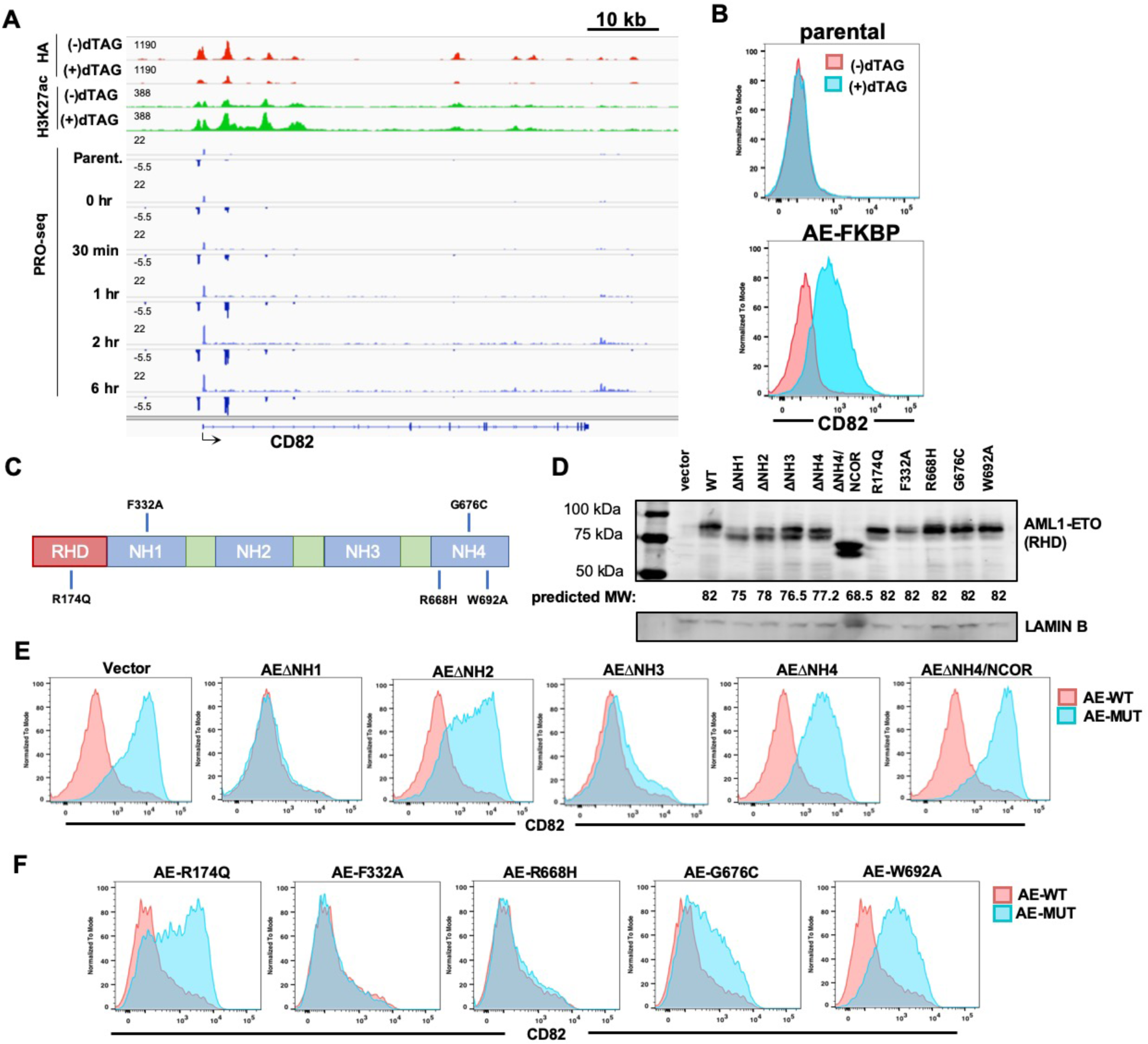
CD82 serves as a reporter for AE-mediated repression. (A) HA Cut&Run, H3K27ac ChIP-seq, and PRO-seq signal are shown at the *CD82* locus before and after AE-FKBP^F36V^-2xHA degradation following dTAG-47 treatment. (B) Flow cytometry for surface CD82 protein shows that dTAG-treatment induces CD82 protein expression in AE-FKBP-expressing cells (lower), but not in parental cells (upper). (C) Schematic illustrates AML1-ETO domain structure and the location of point mutations generated. (D) Western blot shows the expression of wild-type (WT) AML1-ETO and the indicated point mutants or domain deletion mutants. Domain deletion mutations (E) or point mutations (F) in AML1-ETO were expressed in AE-FKBP Kasumi-1 cells in the presence of dTAG-47 to degrade the endogenous AE-protein. CD82 protein levels were quantified by flow cytometry and the ability of each mutant to repress CD82 expression was compared with exogenous wild-type AML1-ETO (AE-WT).

We also performed motif analysis to determine if AML1-ETO bound regions were enriched in other transcription factor binding motifs. Within cluster 1 and 2, the RUNX1 and ETS factor motifs were most frequently observed, which is consistent with RUNX family members cooperatively binding DNA with ETS factors^44,45^. Within the 59 gene signature, RUNX1 and ETS sites again dominated the motif analysis (not shown). Somewhat surprisingly, E protein binding sites were not in the top five in this motif analysis (Supplemental Figure 3B).

Our degron approach is agnostic to the effect of AML1-ETO on gene regulation (i.e., repression vs. activation), as we capture everything using PRO-seq^8,40^. The prior studies that suggested that AML1-ETO was required for gene expression (i.e., activation) used siRNA or shRNAs and RNA-seq was performed between 48 hrs. and 10 days after transfection or infection. For example, *cyclin D2* (*CCND2*) was reported to be a key target for AML1-ETO-mediated activation after siRNA knockdown of AE in Kasumi-1 cells^46^, using our dTAG system, we saw no changes in the rate of transcription of *CCND2* over the 24hr time course in PRO-seq (Supplemental Fig. 3, compare *CCND2* to canonical targets *NFE2* and *RASSF2*). The *CCND2* locus did appear to be bound by AE (although the Cut&Run signal was less robust than at other targets); however, AE degradation did not correlate with a loss of H3K27ac or a loss of active polymerase at this enhancer, suggesting that enhancer activity was not diminished in the absence of AML1-ETO. Because transcriptional regulators such as *NFE2* (Supplemental Fig. 3A), *CEBPA* (Fig. 3E), *PLZF* (Fig. 2H), and *ETO2* (not shown) were activated more than 1.5-fold within 2hr of dTAG-47 addition, it is important to keep in mind that any effects from 4 hours onward could be indirect. For instance, we noted that *CEBPA* levels began to decline by RNA-seq at 24hr after treatment, while *CEBPE* transcription increased (Supplemental Fig. 1E), which is typical of myeloid differentiation transcriptional cascades. Thus, much of the gene activation activity that has been ascribed to AML1-ETO in the past is likely an indirect consequence resulting from activation of secondary transcriptional networks.

### The NH4 domain is required for activity of AML1-ETO

As a further functional test of the AML1-ETO-FKBP allele, we expressed wild type or mutant forms of AML1-ETO in our AML1-ETO-FKBP12 cells and then degrade the endogenous fusion protein to test for complementation. The cell surface marker, *CD82*, is one of the most highly up-regulated members of the 59 gene repression signature (Fig. 4A). Given that FACS analysis for this cell surface protein yielded easily quantifiable results in which CD82 was maximally induced within 24hr of AML1-ETO-FKBP degradation (Fig. 4B), we were able to use surface CD82 expression as a reporter for AE-mediated gene repression. Unexpectedly, a mutant that disrupted with the ability of ETO to interfere with HEB or E2A-mediated transactivation (AE^F332A^)^47,48^ had essentially no impact, as *CD82* was still repressed in the presence of this mutation. Moreover, deletion of the entire NH1 domain, which mediated E-protein binding^47,48^, was fully capable of repressing *CD82* (Fig. 4E). By contrast, ΔNH4, which appears to suppress leukemia development in mice^35^, was unable to fully suppress *CD82* (Fig. 4E), while three point mutations within NH4 (R668H, G676C, or W692A) had varying effects on AE-mediated repression (Fig. 4F). NH4 is responsible for one of the two contacts between NCOR1/SMRT with a second contact site lying between NH1 and NH2 of ETO that is sufficient for co-immunoprecipitation of NCOR1 with AML1-ETO^49-53^. Deletion of the second NCOR1 binding domain in the context of ΔNH4, completely eliminated AML1-ETO-mediated repression of *CD82* (Fig. 4E) and was even more dramatically affected than deletion of the tetramerization domain (ΔNH2, Fig. 4E) or disruption of DNA-binding by the AE^R174Q^ mutation (Fig. 4F).

### Rapid loss of self-renewal and differentiation upon degradation of AML1-ETO in long-term cultures of CD34+ human stem cells

The relatively modest effects on the growth of Kasumi-1 cells upon degradation of AML1-ETO or suppression of AML1-ETO using shRNAs^16,38,39^, suggested that these cell lines may have sustained other mutations or epigenetic changes that allowed them to continue to proliferate in the absence of AML1-ETO, albeit in a more differentiated state (Fig. 1). Mouse models of AML1-ETO-induced leukemogenesis require a truncated protein that is dramatically diminished in its ability to mediate repression (see ΔNH4 and W692A in Fig. 4), and therefore, may not be reflective of the transcriptional networks established by AML1-ETO in human disease^54^. Thus, we turned to long-term cultures of CD34^+^ cells from human cord blood to establish a pre-leukemic system to study AML1-ETO transcription functions. Expression of AML1-ETO in these primary cells impaired myeloid differentiation, even in the context of cytokines that promoted myelopoiesis^23^ to establish a long-term, pre-leukemic state^55^. We expressed AML1-ETO or AML1-ETO-FKBP12^F36V^ in CD34+ HSPCs and found that the addition of the FKBP12 moiety did not affect the ability of AML1-ETO to induce long-term growth of myeloid cultures, or to suppress the expression of *CD82* (Supplemental Fig. 5). In contrast to the results in the established cell lines, when AML1-ETO-FKBP12^F36V^ was degraded by using dTAG-47 (Fig. 5A), not only was CD34 lost from the cell surface (Fig. 5C), but the cells underwent complete myeloid differentiation as indicated by the expression of CD11B, morphological changes, and loss of proliferation (Fig. 5B-E). In contrast, CD34^+^ cultures expressing wild-type AML1-ETO were unaffected by dTAG-47 treatment (Supplemental Fig. 6), suggesting that the differentiation observed in AML1-ETO-FKBP12^F36V^-expressing cells was a direct result of AML1-ETO loss.

**Figure 5.**
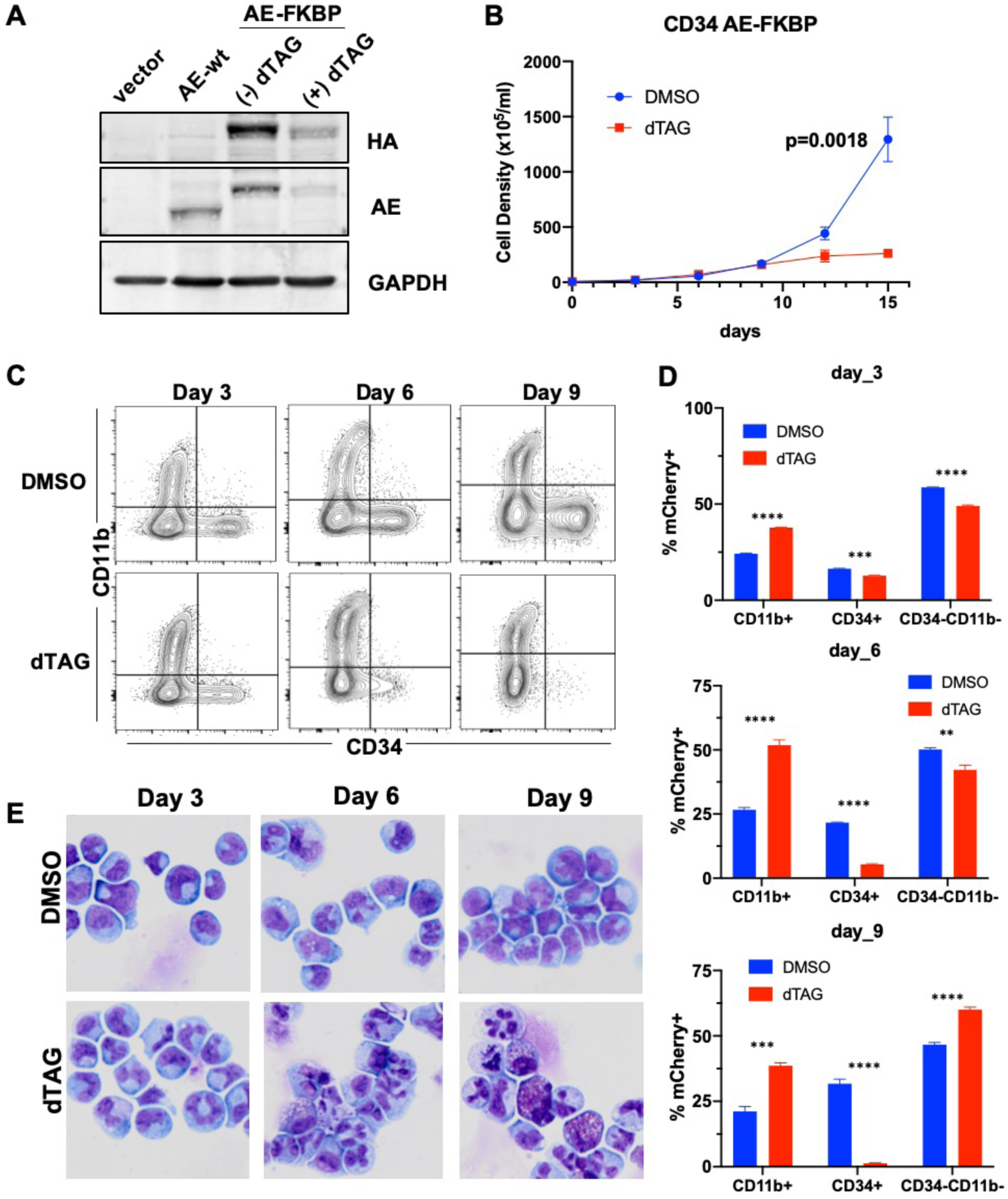
Maintained AE-expression is required to suppress differentiation of CD34^+^ HSPCs. (A) CD34^+^ human cord blood cells were retrovirally transduced with either wild-type AML1-ETO (AE-wt), AML1-ETO-FKBP12^F36V^-2xHA (AE-FKBP), or vector control. AE-FKBP cells were treated with 500 nM dTAG-47 overnight and western blotting of protein lysates used to assess AE protein degradation. (B) Growth curves were performed on CD34 AE-FKBP cells in the presence of 500 nM dTAG-47 or DMSO control. Cells were counted and re-seeded in fresh dTAG-containing media every 3 days for 15 days. (C) AE-FKBP cultures were subject to flow cytometry on 3, 6 and 9-days post-treatment with dTAG-47 to monitor CD34 and CD11b expression. The results of the flow cytometry is quantified in (D). (E) Cytospins were performed every three days on AE-FKBP cultures. Cells were fixed and subject to Wright-Giemsa staining.

Due to the limited number of cells available in this culture system, we used CUT&RUN to ensure that the exogenously expressed AML1-ETO-FKBP12^F36V^ bound to DNA appropriately (Fig. 6A). The majority of robust binding sites overlapped with the CUT&RUN data from Kasumi-1 cells (Fig. 6A and 6B). Next, again due to limited cell numbers, we used RNA-seq at a short interval (4 hours) after addition of dTAG-47 to the culture media to examine changes in gene expression. By comparing these data with the 4 hour RNA-seq data from Kasumi-1 cells, we found that 50 of the 107 genes up-regulated in CD34^+^ cells were induced in both systems. While this concordance is very high, it likely underestimates the similarities, because the short time frame used for RNA-seq to capture direct changes in the CD34+ cells failed to capture canonical targets such as *CD82, OGG1* and *RASSF2*, which were trending upward but did not reach our statistical cutoff within 4 hours. In fact, CD82 protein levels were significantly up-regulated in CD34^+^ cultures by 24 hours post-dTAG treatment (Fig. 6E). Additionally, we observed changes that were unique to the CD34^+^ system such as *GFI1B* and key stemness genes such as *MYCN* (Fig. 6D and 6F).

**Figure 6.**
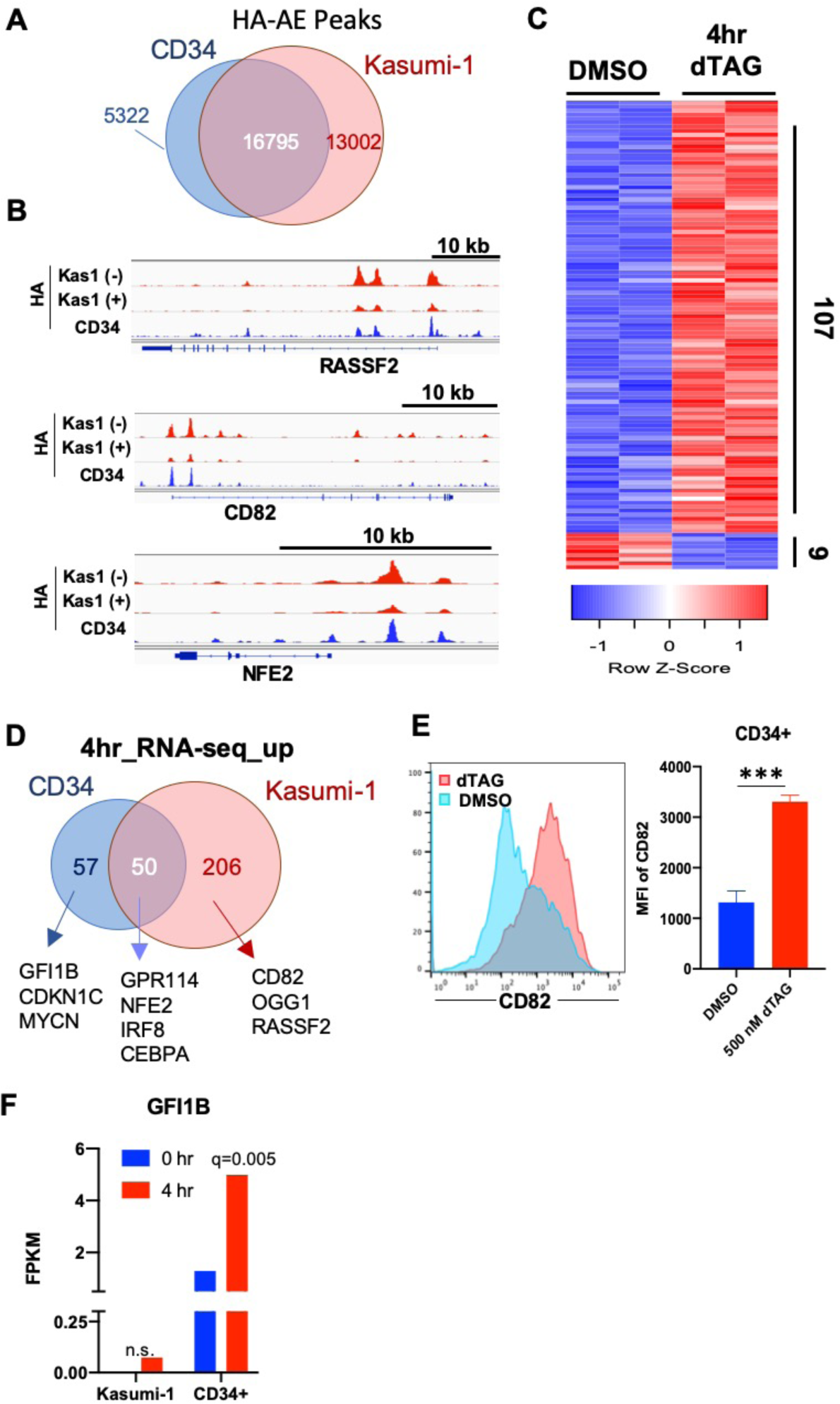
Cut&Run and RNA-seq identify common AE targets in Kasumi-1 cells and primary HSPC cultures. (A) Cut&Run analysis utilizing anti-HA to detect AE-FKBP binding sites in CD34^+^ HSPC cultures. A Venn diagram illustrates the overlap of peaks in Kasumi-1 cells and CD34^+^ HSPC cultures. (B) Screenshots show common AE-binding sites in Kasumi-1 cells and CD34 cultures at the *RASSF2, CD82*, an *NFE2* loci. (C) RNA was isolated from CD34 AE-FKBP cultures 4 hours after dTAG treatment and RNA-seq was performed. The heatmap shows genes significantly changed by greater than 1.5-fold. The 107 genes significantly up-regulated upon AE-FKBP degradation (up > 1.5-fold; q<0.05) were compared with the genes identified by RNA-seq as up-regulated by 4 hours in Kasumi-1 cells (D). (E) Flow cytometry for CD82 shows a significant increase in CD82 protein levels 24 hours following dTAG treatment. (F) RNA-seq revealed a significant increase in *GFI1B* expression following AML1-ETO degradation in CD34 cells, but not Kasumi-1 cells.

GFI1B is particularly intriguing, as it is a RUNX1-regulated gene that is critical for megakaryopoiesis and *RUNX1* mutations are linked to familial platelet disorders that predispose these patients to AML development^56^. While parental Kasumi-1 cells did not fully differentiate in response to degradation of AML1-ETO, LSD1 inhibitors activate *GFI1B*^57-59^. Therefore, we tested whether the combination of degrading AML1-ETO in the context of LSD1 inhibitor (100 nM GSK2879552) with or without all trans-retinoic acid (ATRA) would activate GFI1B and trigger differentiation. The combination of LSD1 inhibitor and degrading AML1-ETO in Kasumi-1 cells caused the firing of an AML1-ETO-bound enhancer 3’ to the *GFI1B* gene within 24 hours (Fig. 7A). Moreover, this drug combination had a dramatic impact on *GFI1B* expression by 3 days (Fig. 7B). While dTAG-47 + LSD1 inhibitor alone induced some differentiation in Kasumi-1 cells, degrading AML1-ETO dramatically sensitized these cells to a combination of ATRA+ LSD1 inhibitor (Fig. 7C, 7D). Thus, even in a cell line that contains additional mutations not commonly observed in t(8;21) patients (e.g., mutant *TP53*), drugging AML1-ETO had important biological and potentially therapeutic effects.

**Figure 7.**
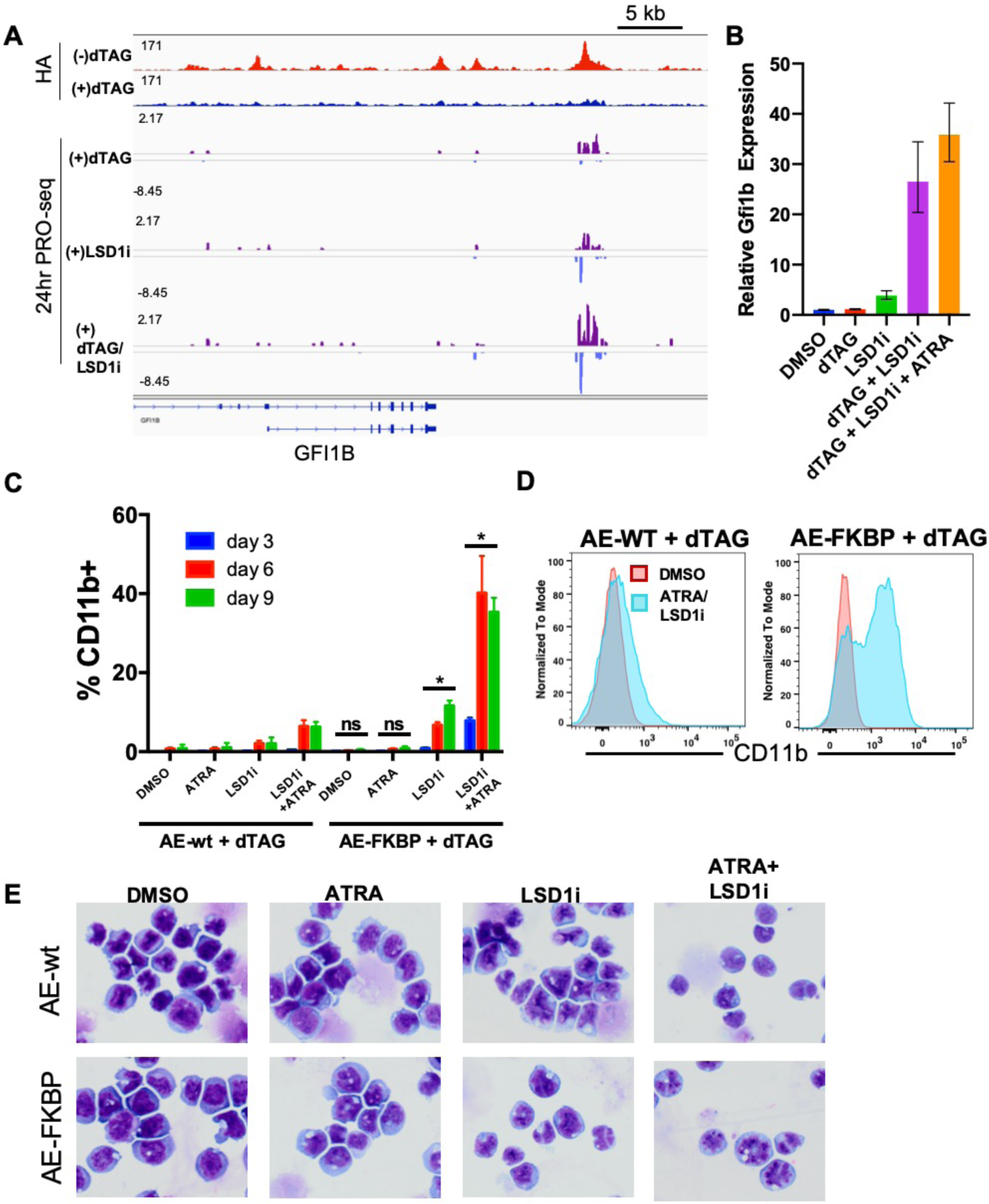
Combining dTAG treatment with LSD1 inhibition and ATRA causes differentiation of Kasumi-1 cells. (A) Cut&Run using anti-HA identified a 3’ enhancer bound by AML1-ETO at the *GFI1B* locus in Kasumi-1 cells. PRO-seq revealed an increase in eRNA transcription at this site following a 24-hour treatment with both dTAG-47 and 100nM of the LSD1 inhibitor, GSK2879552, but not with either molecule alone. (B) Kasumi-1 AE-FKBP cells were treated with dTAG, LSD1i, dTAG and LSD1i, or dTAG + LSD1i +ATRA for three days prior to isolation of RNA. Q-RT-PCR was performed to determine the relative expression of *Gfi1b* compared to *Actb* control. Expression is shown relative to DMSO treated cells and error bars represent the RQ_min_ and RQ_max_. (C) Parental Kasumi-1 cells (AE-wt) or AE-FKBP Kasumi-1 cells were treated with 100 nM dTAG alone or in combination with 100 nM GSK2879552, 1 μM ATRA, or both. At indicated timepoints, CD11b^+^ cell populations were quantified by flow cytometry. (D) Representative flow cytometry plot from 6 days post-treatment. (E) Cytospins were performed 6 days following treatment with dTAG-47 and DMSO, ATRA, LSD1 inhibitor, or ATRA + LSD1i. Following fixation, cells were subject to Wright-Giemsa staining.

## Discussion

One of the biggest challenges in studying transcription factor function is to define the direct, primary targets of the factor. Early studies of DNA binding proteins used foot-printing methods and reporter assays to define whether a factor could activate or repress transcription, but these one-gene-at-a-time approaches were not only slow and laborious, but also limited to *in vitro* DNA binding or transfected plasmids. Thus, the results described what the factor could do in an artificial system, but not how it acted genome-wide and in the context of native chromatin to control gene expression networks. The development of modern genomic methods (e.g. ChIP-seq and RNA-seq) coupled with siRNA approaches allowed genome-wide analysis, yet most DNA binding sites could not be easily assigned a function. This is perhaps not surprising for a transcription factor that binds a six base pair consensus site, as this yields far more potential binding motifs than genes to regulate. Furthermore, waiting two to ten days before measuring gene expression made it difficult to deconvolute direct vs. indirect transcriptional changes. Thus, in many cases, genes that were thought to be direct targets of a transcription factor were the result of phenotypic changes, rather than direct changes in gene expression that drove that phenotype. By integrating a small molecule-responsive degron tag into the endogenous locus of a transcription factor and coupling the rapid degradation of this factor to genome-wide nascent transcription assays, we have established a system to collapse the time frame for transcription factor analysis from days to minutes, which allowed the identification of a core transcriptional network with functional DNA binding sites, while also defining the mechanism of action.

The t(8;21) fusion protein is associated with a 5-year survival rate of only about 50%, underscoring the need for additional therapeutic targets informed by improved mechanistic insight. AML1-ETO was initially shown to repress transcription through the recruitment of histone deacetylase complexes^52,53^, but more recent reports have suggested a potential role of AML1-ETO-dependent transactivation via interaction with p300^17^ or JMJD1C^15^ to drive expression of mediators of cell cycle progression such as *CCND2*^16^. While our studies confirm the ability of AML1-ETO to repress key targets that control myeloid differentiation such as *CEBPA, NFE2, PLZF* and *CBFA2T3* (*ETO2*), there were few consistent changes to support the hypothesis that AML1-ETO plays any appreciable role in gene activation in either Kasumi-1 cells or primary CD34^+^ HSPC cultures.

Chromosomal translocations that create oncogenic fusion proteins are the ideal therapeutic targets, as these new proteins are not expressed in normal cells. While inhibitors of the BCR-ABL kinase are the prototypical example of the impact that such a drug can have, targeting transcription factors created by translocations has been more challenging. The lone success story is PML-RARA, which drives development of acute promyelocytic leukemia. Even before the breakpoint was identified, all trans-retinoic acid (ATRA) was found to cause myeloid differentiation in this disease, and later arsenic trioxide was found to trigger the degradation of the fusion protein. When used in combination, all trans-retinoic acid and arsenic are potent differentiation agents that have changed the natural history of this disease. Like arsenic in their ability to trigger protein degradation, the PROTACs, typified by lenalidomide, can link transcription factors such as IKAROS to the ubiquitin degradation system^33,34^. In this study, we utilized CRISPR-Cas9 genome editing to create a synthetic PROTAC system for the degradation of the AML1-ETO fusion protein in Kasumi-1 cells. While Kasumi-1 cells became less stem-like after degradation of AML1-ETO, they did not fully differentiate. This is perhaps not surprising given the additional mutations and epigenetic changes that these cells acquired throughout leukemia development and perhaps during establishment of the cell line. In contrast, CD34^+^ human cord blood stem and progenitor cells showed the expected impairment of differentiation upon expression of AML1-ETO-FKBP12^F36V^, and fully differentiated upon AML1-ETO degradation (Fig. 6).

One difference between Kasumi-1 cells and CD34^+^ HSPC cultures was the re-expression of *GFI1B* upon AML1-ETO degradation specifically in CD34^+^ cultures. This was in spite of the fact that AML1-ETO was clearly recruited to the 3’ *GFI1b* locus in both CD34^+^ cultures and Kasumi-1 cells. Recent studies have identified GFI1 and GFI1B, two transcription factors that autoregulate their own expression, as critical targets of LSD1 inhibition^57-59^, thus we explored whether combining an LSD1 inhibitor with AML1-ETO degradation could re-activate *GFI1B* expression in Kasumi-1 cells. Not only did this combination result in a dramatic increase in *GFI1b* expression, but it also correlated with an increase of Kasumi-1 cell terminal differentiation (Fig. 7). Moreover, consistent with previous reports of cooperativity between LSD1 inhibitors and ATRA in AML differentiation models^60^, the addition of ATRA to the dTAG/LSD1i combination further facilitated the differentiation of Kasumi-1 cells. Thus, the targeting of AML1-ETO may have therapeutic benefit in combination with differentiation therapy in t(8;21) AMLs.

Finally, by combining AML1-ETO protein degradation with nascent transcript analysis by PRO-seq, we have defined the mechanism of action of the fusion protein, as well as a small core transcriptional signature. By identifying the direct transcriptional targets, these genes can be used as biomarkers for an eventual clinical trial of AML1-ETO-targeted drugs. A prime example of this was CD82, the expression of which varied widely in the presence and absence of AML1-ETO. In fact, flow cytometry analysis of CD82 was so sensitive, that it could be used to determine the relative repression achieved by numerous AML1-ETO point mutants and domain deletions (Fig. 4). Thus, this provides a simple flow cytometry-based assay to quantify the on-target effects of any future AML1-ETO targeted therapies.

## Acknowledgements

We thank the members of the Hiebert lab for helpful discussions, reagents and advice. We thank the Flow Cytometry and Genome Sciences (VANTAGE) Shared Resources for services and support. The content is solely the responsibility of the authors and does not necessarily represent the official views of the NIH. This work was supported by the T. J. Martell Foundation, the Robert J. Kleberg, Jr. and Helen C. Kleberg Foundation, National Institutes of Health grants (RO1-CA178030, RO1-CA164605 and R01-CA64140) and core services performed through Vanderbilt Digestive Disease Research grant (NIDDK P30DK58404) and the Vanderbilt-Ingram Cancer Center support grant (NCI P30CA68485). Kristy Stengel was supported by 5 T32 CA009582-26 and a postdoctoral fellowship (PF-13-303-01-DMC) from the American Cancer Society. The project described was also supported by the National Center for Research Resources, Grant UL1 RR024975-01, and is now at the National Center for Advancing Translational Sciences, Grant 2 UL1 TR000445-06. Scott Hiebert received research funding from Incyte Inc. through the Vanderbilt-Incyte Alliance, but these funds did not support this work.

## Figure Legends

**Supplemental Figure 1.** (A) Heatmaps include all genes significantly changed in at least one timepoint and depict the PRO-seq signal for the indicated timepoint relative to 0 hour control samples. (B) Histograms represent the PRO-seq signal around the transcription start site for genes with up-regulated gene body transcription at at least one timepoint. (C) Venn diagrams show the overlap of the 59 gene AE-repression signature identified by PRO-seq with genes significantly changed by RNA-seq at 2, 4, or 8 hours. (D) Venn diagrams illustrate the overlap between the 7 genes down-regulated at all timepoints in PRO-seq data and genes identified as significantly down-regulated by RNA-seq at 2, 4, and 8 hours. (E) RNA-seq FPKM for *CEBPA* and *CEBPE* over time.

**Supplemental Figure 2.** (A) HA Cut&Run analysis identified over 30,000 HA-AE peaks in Kasumi-1 AE-FKBP cells. Diffbind was used to assess MACS2 called peaks for significant change following dTAG treatment. Significantly changed peaks are designated in red. (B) Cut&Run for wild-type AML1-ETO from parental Kasumi-1 cells and Cut&Run for HA from AE-FKBP12^F36V^-2xHA cells were compared with published ChIP-seq data for AML1-ETO from Kasumi-1 cells^44^. The venn diagram depicts the overlap of MACS2 called peaks for AE ChIP-seq, AE Cut&Run and differential HA peaks (minus dTAG vs. plus dTAG) identified by DiffBind. (C) Heatmaps show the indicated ChIP or Cut&Run signal plotted around all 24,258 called AE ChIP-seq peaks from the previously published data set^44^.

**Supplemental Figure 3.** (A) Pie charts show the number of HA-AE peaks associated with the 59 gene AE repression signature relative to the total number of peaks per cluster. (B) Motif analysis identified the most enriched transcription factor binding motifs in each of the three HA-AE clusters.

**Supplemental Figure 4.** Screenshots depict changes in HA and H3K27ac 24 hours post-treatment with dTAG-47 and changes in PRO-seq signal at the indicated timepoints following dTAG treatment at the *NFE2* (A), *RASSF2* (B), and *CCND2* (C) loci.

**Supplemental Figure 5.** (A) Percentage of mCherry+ cells were monitored over time in MSCV (vector), wild-type AML1-ETO (AE), or AML1-ETO-FKBP12^F36V^-2xHA (AE-FKBP) transduced CD34^+^ human cord blood cultures. Percentage of mCherry relative to week 1 post-infection is plotted over time. (B) Flow cytometry was performed at week 4 post-infection to determine the relative percentage of CD34^+^ vs. CD11b^+^ cells present in HSPC cultures. Quantification of three independent infections is displayed in the graph on the right. (C) Flow cytometry for cell surface CD82. Quantification at the right shows the average mean fluorescence intensity (MFI) from at least three independent cultures.

**Supplemental Figure 6.** (A) Parental Kasumi-1 cells (AE-WT) were counted every three days for 15 days following treatment with 500 nM dTAG-47. (B) Parental Kasumi-1 cells were monitored for relative CD34 and CD11b expression at 3, 6, and 9 days post-dTAG treatment. (C) Cytospins of dTAG-treated parental Kasumi-1 cells were subject to Wright-Giemsa staining.

